# What is behavior? No seriously, what is it?

**DOI:** 10.1101/2021.07.04.451053

**Authors:** Adam Calhoun, Ahmed El Hady

**Author notes:** correspondence should be addressed to Ahmed El Hady or Adam Calhoun.

## Abstract

Studying ‘behavior’ lies at the heart of many disciplines. Nevertheless, academics rarely provide an explicit definition of what ‘behavior’ actually is. What range of definitions do people use, and how does that vary across disciplines? To answer these questions we have developed a survey to probe what constitutes ‘behavior’. We find that academics adopt different definitions of behavior according to their academic discipline, animal model that they work with, and level of academic seniority. Using hierarchical clustering, we identify at least six distinct types of ‘behavior’ which are used in seven distinct operational archetypes of ‘behavior’. Individual respondents have clear consistent definitions of behavior, but these definitions are not consistent across the population. Our study is a call for academics to clarify what they mean by ‘behavior’ wherever they study it, with the hope that this will foster interdisciplinary studies that will improve our understanding of behavioral phenomena.

## Introduction

The scientific definition of ‘behavior’ has tended toward the maxim ‘I know it when I see it’. But does everyone know and see the same thing? Much ink has been spilled coming up with precise definitions of behavior, stretching from Aristotle [1], through Wiener’s cybernetics [25], Von Neumann [23], Lorenzian ethology [13], Tinbergen behavioral theory [20] and Von Uekuell’s umwelt [21]. More recent work has operationalized such definitions to create machine learning algorithms capable of ‘recognizing’ when an animal is performing some ‘behavior’ [2, 26, 3]. An alternative to creating one’s own definition has been to ask how experts in a field actually use the word. Within behavioral biology, Levitis et al [12] surveyed the literature and found 25 distinct operational definitions of behavior. These range from the vague (‘what an animal does’, [4]) to the specific (‘A response to external and internal stimuli, following integration of sensory, neural, endocrine, and effector components). Surveys on biological concepts such as ‘genes’ and ‘consciousness’ [7, 19] have similarly identified highly divergent meanings of words that form the basis of their respective fields. However, ‘behavior’ is a concept not only used by behavioral biologists but across many fields. To date there are no surveys that cover a wide variety of academic fields when it comes to the definition of behavior.

There are good reasons to believe that different fields use different definitions and that these definitions in turn influence what is studied. For instance, consider the study of a risky choice. The behavior that an economist might be interested in are the biases and heuristics that people employ while performing real world decisions, while a neuroscientist seeks to understand how the ‘behavior’ is mediated via neural mechanisms. On the other hand, behavioral ecologists seek to study how ‘behavior’ is shaped by the environment, while sociologists seek to understand society through the ‘behavior’ of its individuals. While each of these researchers seeks to understand ‘behavior’, they are attempting to understand fundamentally different things by virtue of these disciplinary differences in mechanism,.

This can reflect deeper, more fundamental disagreement about what ‘behavior’ is. For instance, sensorimotor behavior is behavior that relates sensory input to motor output. But is this ‘behavior’ when it is performed by a rat? How about a C. elegans, with only three or four synapses between sensory neurons and motor output? What about a bacteria or a computer program? Are the neurons themselves ‘behaving’? Furthermore, not only is the animal moving but there are cognitive variables and needs that influence the animal’s action. Which of these constitute the ‘behavior’?

Instead of creating our own definition of behavior, we aimed to identify how the word ‘behavior’ is actually used by the academic community. We designed a survey aimed at determining the answer to this question. Through a quantitative analysis of survey data, we show that what constitutes ‘behavior’ differs between academic disciplines, level of academic seniority, and type of animal model organism studied. We further show that there are at least six ‘types’ of behavior that are used to construct seven archetypes of behaviors.

The study pinpoints to the lack of a common inter-disciplinary definition of behavior. We argue that we do not need an agreed-upon singular definition of behavior, but we do have to be clear on what the definitions that we are using are in any given study. Those clear working definitions will open up the opportunity for effective interdisciplinary studies that tackle multiple facets of ‘behavior’.

## Results

### Widespread disagreement of definitions of behavior

We developed a set of questions designed to probe how people define ‘behavior’. For instance, ‘A behavior is always the output of motor activity’, ‘Is learning a behavior?’, and ‘Is a dog that follows a scent trail behaving?’. We included questions that were designed to elicit inconsistencies such as, ‘Is a reflex a behavior?’ and ‘Is sweating a behavior?’. The order of questions was randomized and the survey was distributed online and elicited responses from a range of academic disciplines and seniorities (see Methods, Table 1 and 2). Analysis revealed that respondents did not provide a consistent answer for the the majority of questions: 43 / 48 answers had fewer than 80% of respondents answering in the same manner (Table 1, Supplemental Fig 1a). While there were some questions that showed widespread agreement in either a ‘Yes’ (’Is a dog that follows a scent trail behaving’) or a ‘No’ (’Does a behavior need to be intentional’) answer (Supplemental Fig 1b-d), most questions showed disagreement. The questions with the most disagreement (defined as the highest uncertainty, or entropy, in their responses), included ‘Can computer programs behave?’ and ‘A behavior always involves interaction with the environment?’ (Fig 1a). We remain agnostic as to the source of disagreement between responses: it is possible that different subjects are responding to different aspects of each question. This does not impact the interpretation of our results as we believe this itself provides a consistent signal about underlying beliefs (see Discussion).

**Table 1:** Survey questions

| Question |  | Yes<br>(%) | No<br>(%) | Maybe<br>(%) |
| --- | --- | --- | --- | --- |
| Q0 | Is behavior the output of a sequence of non-motor signals (eg thinking)? | 43.74 | 17.36 | 38.9 |
| Q1 | All behaviors an animal performs can potentially be identified from recorded video data by using a smart enough computer algorithm. | 23.74 | 47.47 | 28.79 |
| Q2 | A person watches a movie. Is that person producing a behavior? | 73.85 | 9.23 | 16.92 |
| Q3 | There is always a precise time you can point to a behavior starting or ending. | 14.29 | 60.22 | 25.49 |
| Q4 | Is a baby urinating a behavior? | 69.01 | 15.16 | 15.82 |
| Q5 | We have to understand what it is like to be a bat to understand its behavior. | 24.4 | 49.45 | 26.15 |
| Q6 | Working memory is temporarily holding something in memory. Is working memory a behavior? | 20.88 | 61.76 | 17.36 |
| Q7 | Behaviors cannot be studied unless anthropomorphized so that the human experimenter can talk about them. | 7.69 | 73.63 | 18.68 |
| Q8 | A person sees food that they like but choose not to move toward it. Is that person behaving? | 84.62 | 7.47 | 7.91 |
| Q9 | Is a reflex a behavior? | 59.78 | 18.46 | 21.76 |
| Q10 | An animal watches a movie and tries to decide if objects are moving left or right. As it sits there, trying to decide about which way objects are moving, is it behaving? | 61.54 | 22.2 | 16.26 |
| Q11 | A person sweats in response to hot air. Is this person behaving? | 28.57 | 57.8 | 13.63 |
| Q12 | There are behaviors that you cannot directly observe from possible movements. | 71.65 | 11.21 | 17.14 |
| Q13 | An animal hears one sound, then another. In its mind, it compares the two sounds. Is it behaving? | 39.34 | 41.32 | 19.34 |
| Q14 | The sucking reflex is when babies instinctively suck anything that touches the roof of their mouth. Is the sucking reflex a behavior? | 72.09 | 14.73 | 13.19 |
| Q15 | Can invertebrates behave? | 90.55 | 0.44 | 9.01 |
| Q16 | A neuron spikes. It produces more spikes when the animal's head is in a particular direction. Is that neuron producing a behavior? | 17.8 | 46.59 | 35.6 |
| Q17 | Is a dog that follows a scent trail behaving? | 95.6 | 1.1 | 3.3 |
| Q18 | Can computer programs behave? | 36.26 | 27.03 | 36.7 |
| Q19 | Is sweating a behavior? | 32.31 | 48.57 | 19.12 |
| Q20 | What behavior an animal is producing can only be understood by considering its unique way of sensing and experiencing the world. | 49.23 | 21.54 | 29.23 |
| Q21 | Is a mouse that is held in place while choosing between two options performing the same behavior as a mouse that is free to move around while choosing between the same two options? | 6.37 | 69.01 | 24.62 |
| Q22 | Does a behavior need to be intentional (does every behavior an animal produces have a purpose)? | 13.85 | 76.26 | 9.89 |
| Q23 | A behavior is always potentially measurable | 56.7 | 15.82 | 27.47 |
| Q24 | Does a behavior always relate to something (eg, walking toward something versus 'just walking')? | 16.7 | 72.75 | 10.55 |
| Q25 | The knee-jerk reflex is when a tap of a hammer results in the leg extending once before coming to rest. Is the knee-jerk reflex in adults a behavior? | 53.63 | 36.26 | 10.11 |
| Q26 | Is behavior the output of a sequence of motor signals (eg walking)? | 41.1 | 15.6 | 43.3 |
| Q27 | When fish are schooling (moving in a group), is the school (the whole group of fish) behaving? | 70.99 | 11.21 | 17.8 |
| Q28 | Is learning a behavior? | 50.55 | 24.18 | 25.27 |
| Q29 | A behavior is always in response to the environment | 18.46 | 65.05 | 16.48 |
| Q30 | Is an adult urinating a behavior? | 78.46 | 6.59 | 14.95 |
| Q31 | You are wearing a virtual reality headset. You see a virtual tree and reach out to touch it. Is this the same behavior as reaching out to a real tree? | 28.57 | 34.73 | 36.7 |
| Q32 | A person sees a video on TV. Is that person producing a behavior? | 51.21 | 17.58 | 31.21 |
| Q33 | An animal is thirsty. It must choose between two levers, one of which will provide water. Would you refer to the animal seeking relief from thirst as the primary behavior the animal was producing? | 42.2 | 28.13 | 29.67 |
| Q34 | Behaviors are always discrete; you are either performing that behavior or you are not. | 21.76 | 56.7 | 21.54 |
| Q35 | A rat has a dislike for salty food. Is disliking salty food a behavior? | 15.82 | 72.09 | 12.09 |
| Q36 | A thermostat turns on air conditioning in response to heat. Is the thermostat behaving? | 33.19 | 49.89 | 16.92 |
| Q37 | Can cells behave? | 52.09 | 23.3 | 24.62 |
| Q38 | A behavior always involves interaction with the environment ? | 32.31 | 44.84 | 22.86 |
| Q39 | Behaviors have magnitude; speaking loudly is not the same behavior as speaking quietly. | 67.25 | 12.97 | 19.78 |
| Q40 | An animal reacts defensively to a picture of a predator. Over repeated exposures to this picture, it slowly stops reacting. Is this adaptation a behavior? | 61.1 | 25.27 | 13.63 |
| Q41 | A bacteria can sense chemicals in its environment. It will turn when it senses more of one type of chemical, and move forward if it senses less of that chemical. Is this bacteria behaving? | 72.31 | 10.77 | 16.92 |
| Q42 | Particles move around. Are the particles behaving? | 18.02 | 65.05 | 16.92 |
| Q43 | Bacteria will release chemicals that attract other bacteria. The bacteria then move around as a group. Are these bacteria producing behavior? | 69.89 | 10.77 | 19.34 |
| Q44 | A behavior is always the output of motor activity | 20.22 | 61.1 | 18.68 |
| Q45 | A person reads a book. Is that person producing a behavior? | 88.35 | 5.05 | 6.59 |
| Q46 | Is a sponge filtering water a behavior? | 28.13 | 44.18 | 27.69 |
| Q47 | Can invertebrates weigh costs and benefits in order to produce behavior? | 52.31 | 5.27 | 42.42 |

**Table 2:** Survey metadata

| Demographic data | N | % |
| --- | --- | --- |
| Seniority | 455 |  |
| professor | 115 | 25.27 |
| postdoc | 148 | 32.53 |
| grad student | 142 | 31.21 |
| undergraduate | 18 | 3.96 |
| Gender | 455 |  |
| Male | 285 | 62.64 |
| Female | 159 | 34.95 |
| Other | 5 | 1.1 |
| Choose not to say | 6 | 1.32 |
| Academic fields | 455 |  |
| Neuroscience | 364 | 80.0 |
| Psychology | 79 | 17.36 |
| Biology | 106 | 23.3 |
| Medicine | 17 | 3.74 |
| Engineering | 28 | 6.15 |
| Mathematics | 21 | 4.62 |
| Statistics | 18 | 3.96 |
| Machine Learning | 39 | 8.57 |
| Ethology | 26 | 5.71 |
| Ecology | 13 | 2.86 |
| Philosophy | 9 | 1.98 |
| Sociology | 13 | 2.86 |
| History | 5 | 1.1 |
| Languages and Literature | 3 | 0.66 |
| Model organism | 455 |  |
| humans | 130 | 28.57 |
| rodents | 160 | 35.16 |
| other mammals | 6 | 1.32 |
| fish | 15 | 3.3 |
| birds | 9 | 1.98 |
| other non-mammalian vertebrates | 5 | 1.1 |
| drosophila | 16 | 3.52 |
| c. elegans | 13 | 2.86 |
| other invertebrates | 16 | 3.52 |
| in silico | 35 | 7.69 |

**Figure 1:**
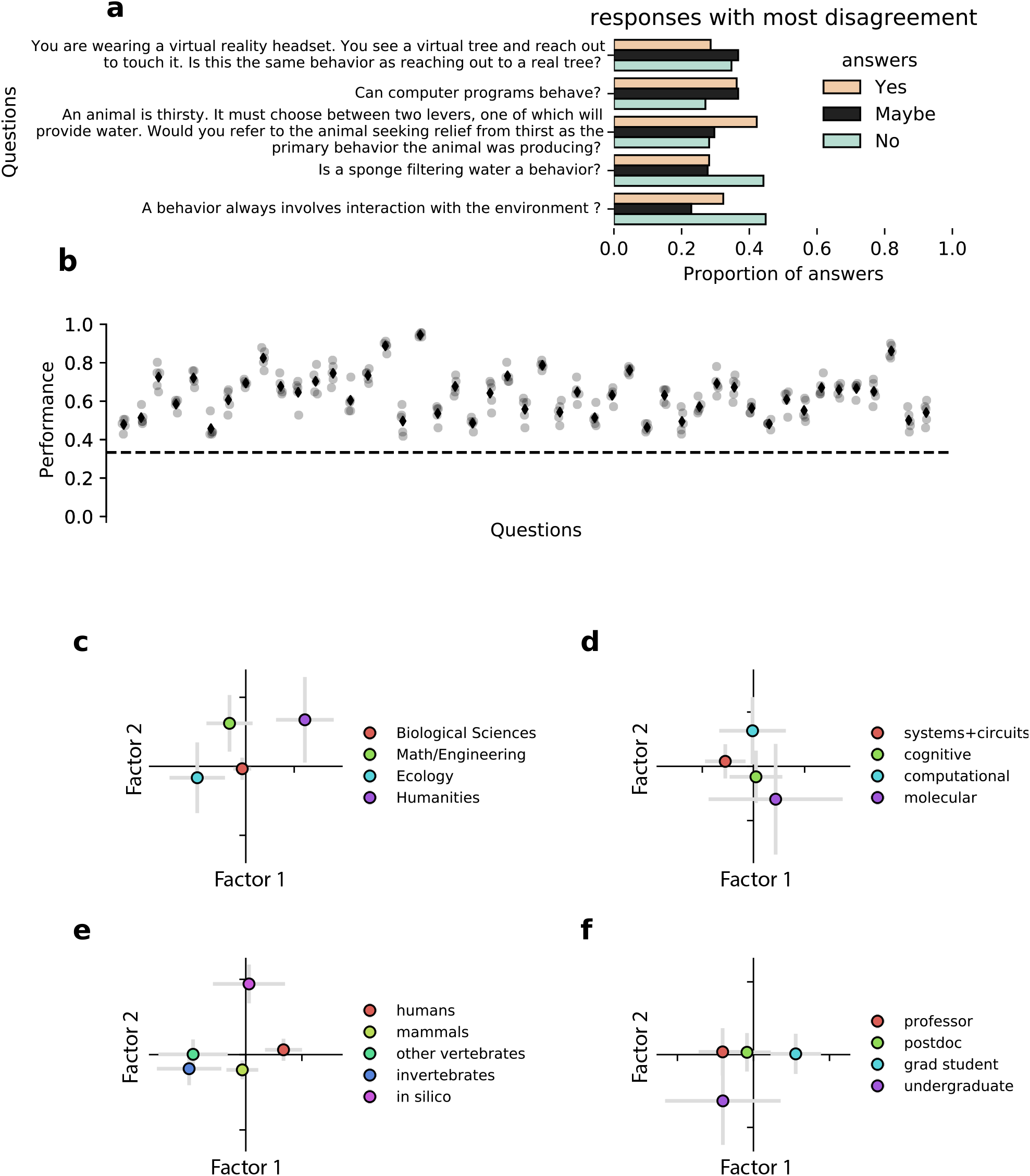
Definitions of behavior are varied but internally consistent. **a**, The five questions with the most disagreement between respondents as defined by the response entropy (n=455). **b**, Responses are predictable with a regression model. Gray dots represent the performance at predicting answers on held-out data using 5-fold cross-validation, black diamond is mean. Mean prediction accuracy across responses is 63%. The dashed line represents chance performance. **c-f**, We used a factor analysis (Multiple Correspondence Analysis (MCA)) to reveal how groups responded on similar questions. Plotting responses on the two largest factors, colored by (**c**) scientific field, (**d**) neuroscience specialty, (**e**) model organism used, and (**f**) academic seniority. All data shown is the mean, and error bars are SEM.

A previous survey of professionals in animal behavior[12] had observed that respondents were inconsistent and self-contradictory. Was this true for our survey? We used a regression model to ask whether we could predict responses from the answers to previous questions and found that we could (Fig 1b). This was true both on average and across all questions. This suggests that disagreement on behavioral questions represents disagreement on the underlying definition of behavior itself.

We then asked whether this variation in responses could be due to differences in academic disciplines - perhaps those trained in the Humanities would have a fundamentally different idea of ‘behavior’ from those trained in Engineering, for instance. We embedded responses in a lower-dimensional space using multiple correspondence analysis (MCA), a factor analysis technique for categorical variables [11] (Supp Fig 2). For visualization purposes, we grouped subjects into broad academic categories: Biological Sciences, Math/Engineering, Ecology, and Humanities (see Methods for definitions, but see Supplemental Fig 3 for all disciplines). We found differences between each field (Fig 1c, Supp Fig 3a, 4a). We hypothesized that even within a field there might be substantial differences; a Cognitive Neuroscientist gets different training from a Systems Neuroscientist. Again, plotting the mean responses for each sub-field showed significant differences (Fig 1d, Supp Fig 4b).

In behavioral research, scientists can ask fundamentally different questions about behavior when they work with humans than when they work with invertebrates. For instance, C. elegans researchers typically investigate chemotaxis and thermotaxis, which are activities that involve sensorimotor transformations, while researchers who study humans may be more likely to study cognitive activities like abstract decision-making. Additionally, those who study invertebrates can map behavior to a single-neuron level, something not typically done in humans. We thus divided subjects by the model organism that they work with. We again find differences between model organisms (Fig 1e, Supp Fig 3b,4c). Surprisingly, we also found that academic experience is also associated with differences in behavioral definitions (Fig 1f, Supp Fig 4d).

### Hierarchical clustering finds definitions

One possibility is that these differences between fields are not because each has a different single definition of behavior, but because certain behavioral definitions are more common in that field. In order to identify potential definitions of behavior, we performed hierarchical clustering on both the responses and respondents (Fig 2). We found six potential classes of questions and seven potential classes of responses.

**Figure 2:**
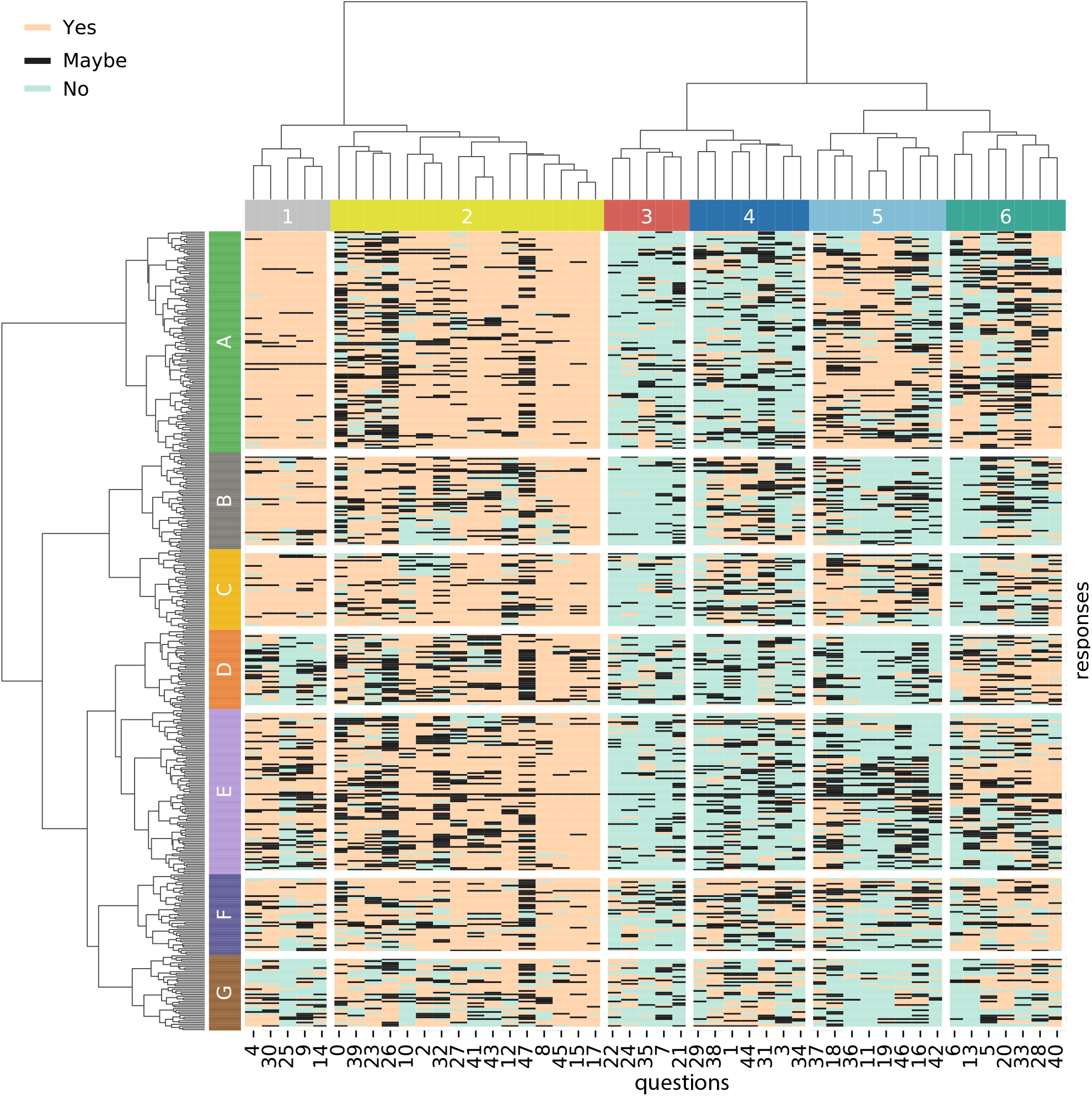
Survey responses reveal clustered definitions of behavior. Hierarchical clustering on the 455 responses reveals six clusters in the questions and seven clusters in the responses. The six categories of questions, labeled 1-6, are particular types of behavior, and the seven categories of responses, labeled A-G, are behavioral definitions. See Figures 3 – 4.

We began by identifying whether the different clusters were meaningful. Examining the questions in the response clusters revealed that similar questions were in each cluster (Fig 3). For instance, Cluster 1 was filled with questions about whether reflexes count as behaviors (’Is a baby urinating a behavior?’, ‘Is a reflex a behavior?’, ‘Is the sucking reflex a behavior?’, and so on). Based on this, we term these ‘behavior definition clusters’ and labeled them as:

1. Reflex (Fig 3a)
2. Actions (Fig 3b)
3. Understanding the mind (Fig 3c)
4. Motor or sensorimotor (Fig 3d)
5. Non-animal (Fig 3e)
6. Cognition (Fig 3f)

**Figure 3:**
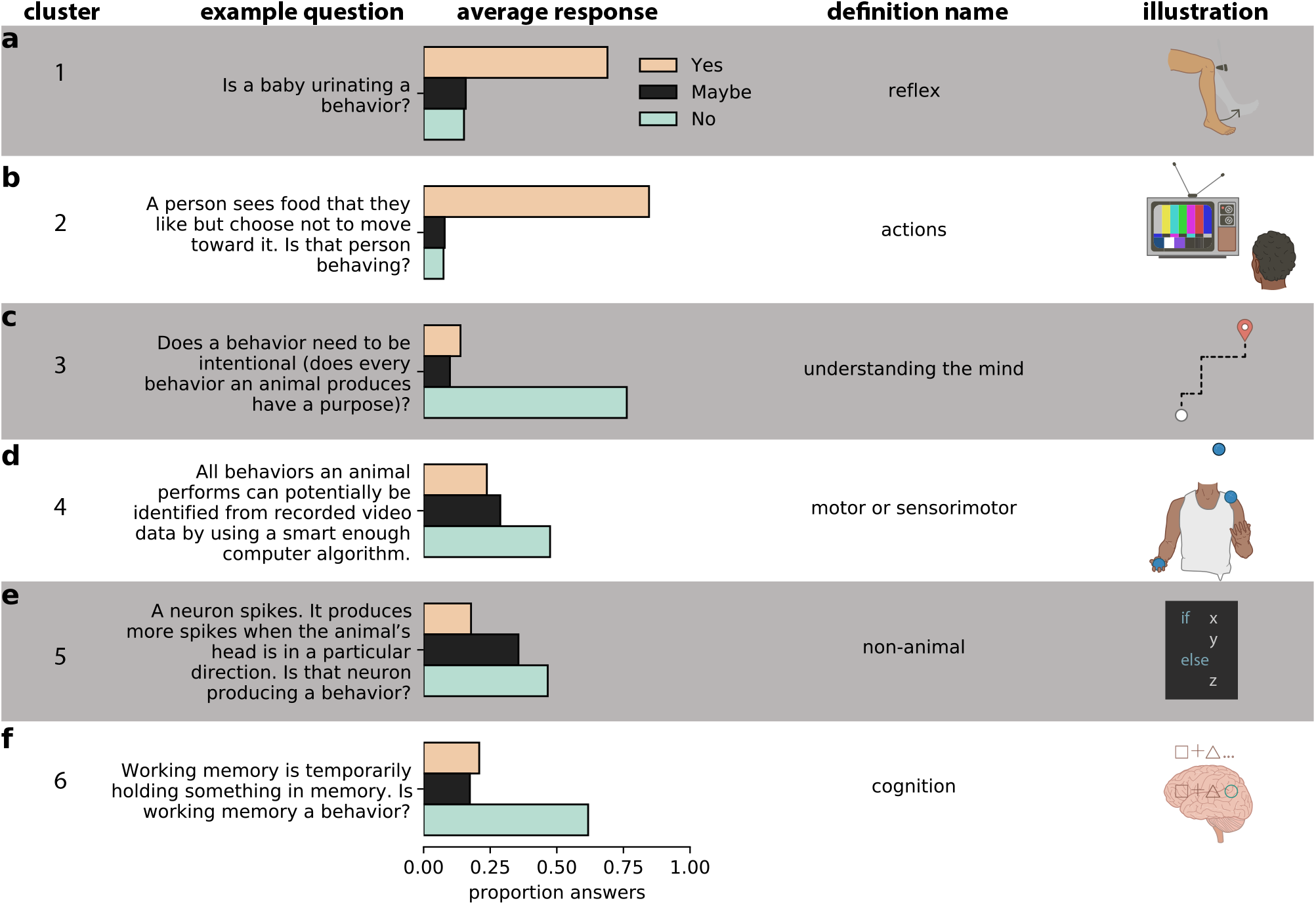
Behavioral archetypes revealed by clustering. **a-f**, Example questions for each type of behavior (left). Illustration representing that type of behavior (right). These definitions are (a) Reflex, (b) actions are behavior, (c) we must understand the mind of an animal to identify its behavior, (d) motor or sensorimotor, (e) non-animal behaviors, (f) learning and memory/cognition.

### Behavioral archetypes

We next examined what each response cluster corresponds to. For each respondent cluster - which we call a ‘behavioral archetype’, answers were relatively consistent in each behavioral definition but answers varied widely between clusters (Fig 4a, Supplemental Fig 5). For instance, in the ‘reflex’ behavioral definition the mean across all responses was 67% ‘Yes’. However, the ‘A’ archetype had 94% of all responses as ‘Yes’ while the ‘D’ and ‘G’ clusters had 19% and 25% of responses as Yes, respectively. To better understand how each archetype corresponds to the respondent’s answers, we plotted the change in responses relative to the overall mean for ‘Yes’, ‘No’, and ‘Maybe’ responses (Fig 4b, Supp Fig 6).

**Figure 4:**
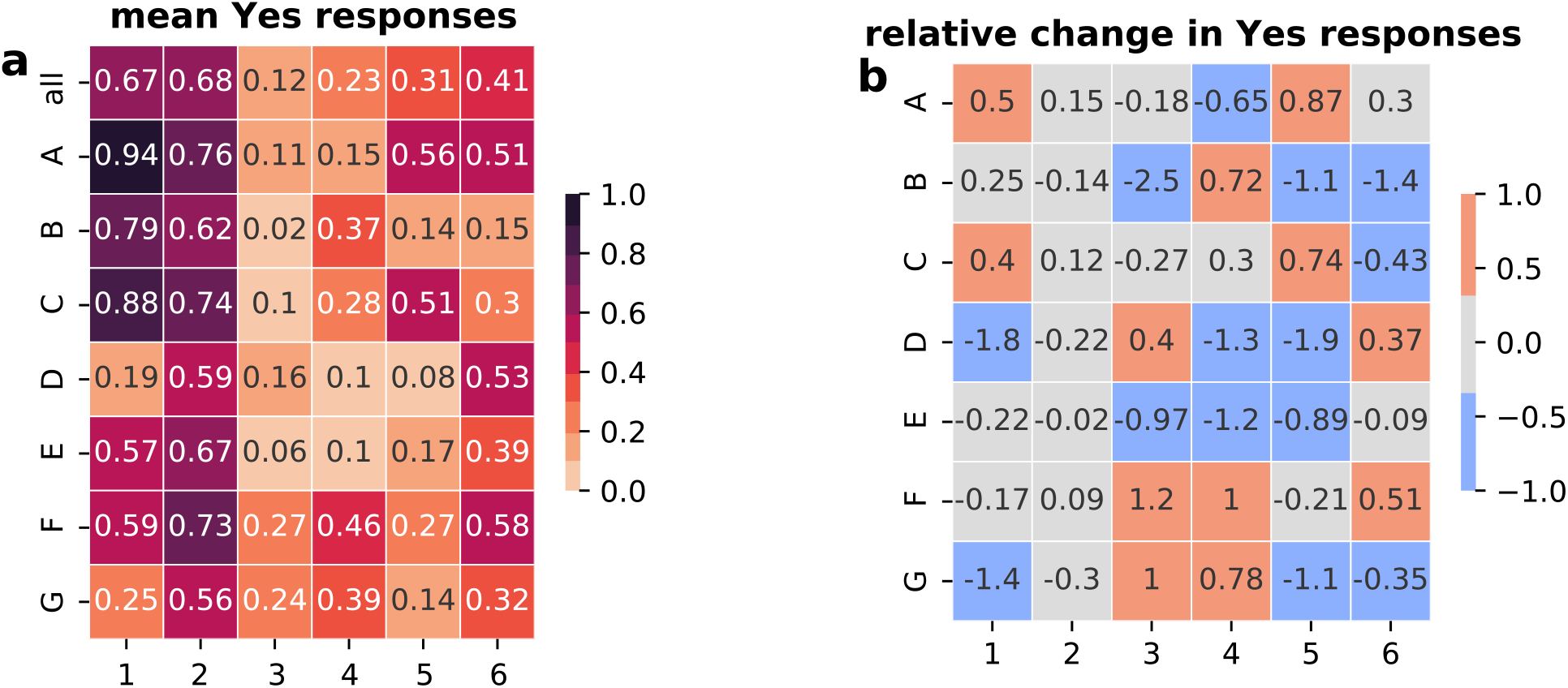
Validation and consistency of behavioral archetypes. **a** Responses are consistent (regression). **b**, The mean ‘Yes’ response in each category allow us to understand what the definitions are composed of.

Based on these, we named these archetypes by examining which behavioral definitions were being used:

A. Broad
B. Motor outputs (animals only)
C. Broad (but mostly motor)
D. Cognition (and anthropomorphizable animal behaviors)
E. Broad (animals only)
F. Well-defined/understood animal behaviors
G. Animals acting with intentions

Taking this all together, we show how behavioral archetypes are made up of different behavioral definitions (Fig 5).

**Figure 5:**
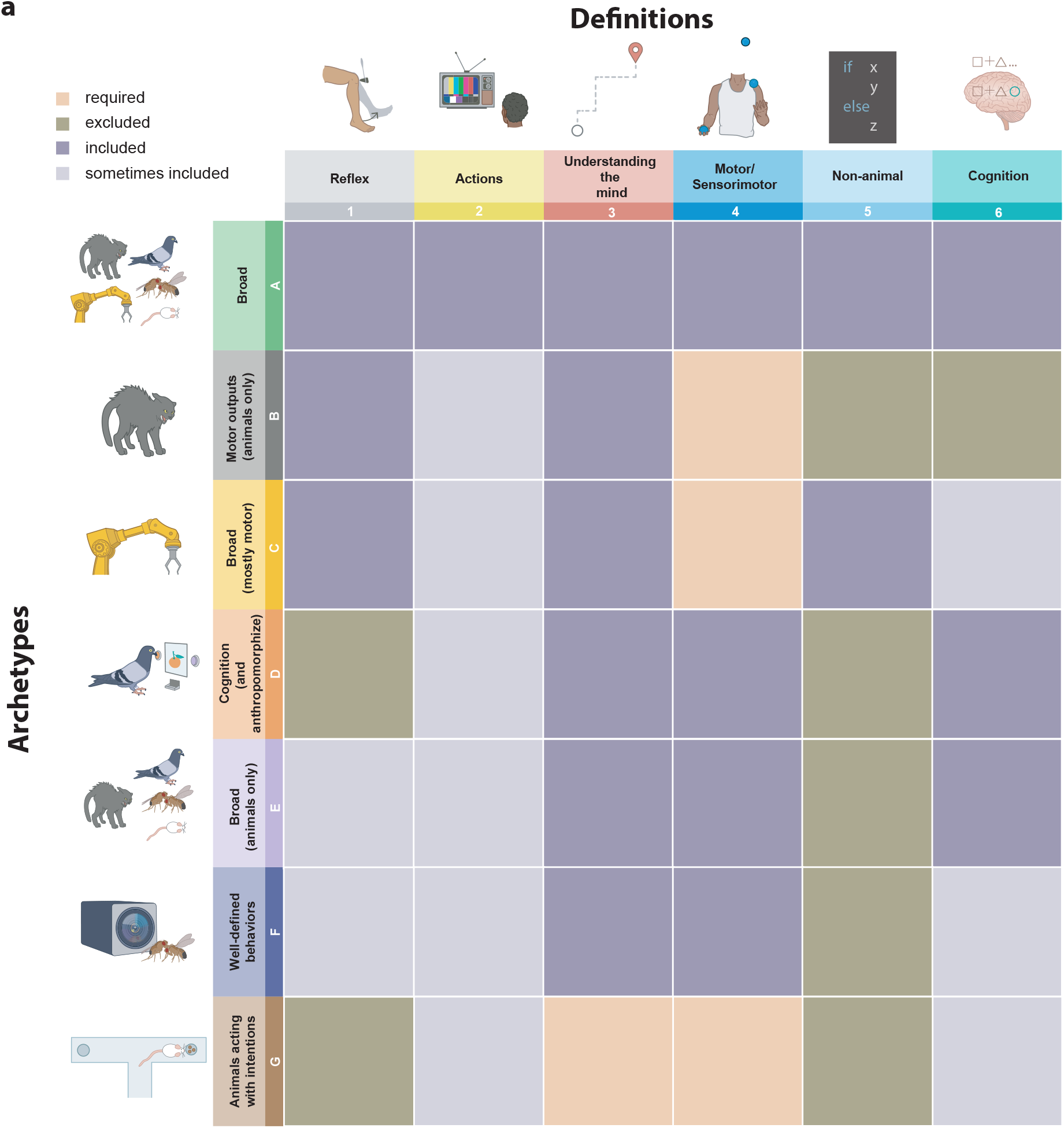
Validation and consistency of behavioral archetypes. **a**, Behavioral archetypes are built out of the responses to different definitions. To illustrate the archetypes, we use the relative responses (Fig 4, Supp Fig 5–6). For each archetype, we color the definitions by whether they are required (pale pink), excluded (brown), included (purple), or sometimes included (grey).

### Academic fields use different definitions

We next used these behavioral archetypes to ask whether different academic fields are consistently using different archetypes when they refer to ‘behavior’. Respondents who come from the humanities were dramatically less likely to be part of the ‘broad’ (A) archetype and much more likely to be a part of the ‘acting with intentions’ (G) archetype. Ecologists were relatively more likely to be part of the ‘broad (mostly motor)’ (C) and ‘broad (animals only)’ (E) archetypes (Fig 6a). Within neuroscience specialties, molecular neuroscientists were more likely to be a part of the ‘broad (mostly motor)’ (C) archetype and cognitive to be part of the ‘well-defined/understood behaviors’ (F) archetype. The ‘broad (animals only)’ (E) archetype was almost entirely Systems and Computational neuroscientists (Fig 6b).

**Figure 6:**
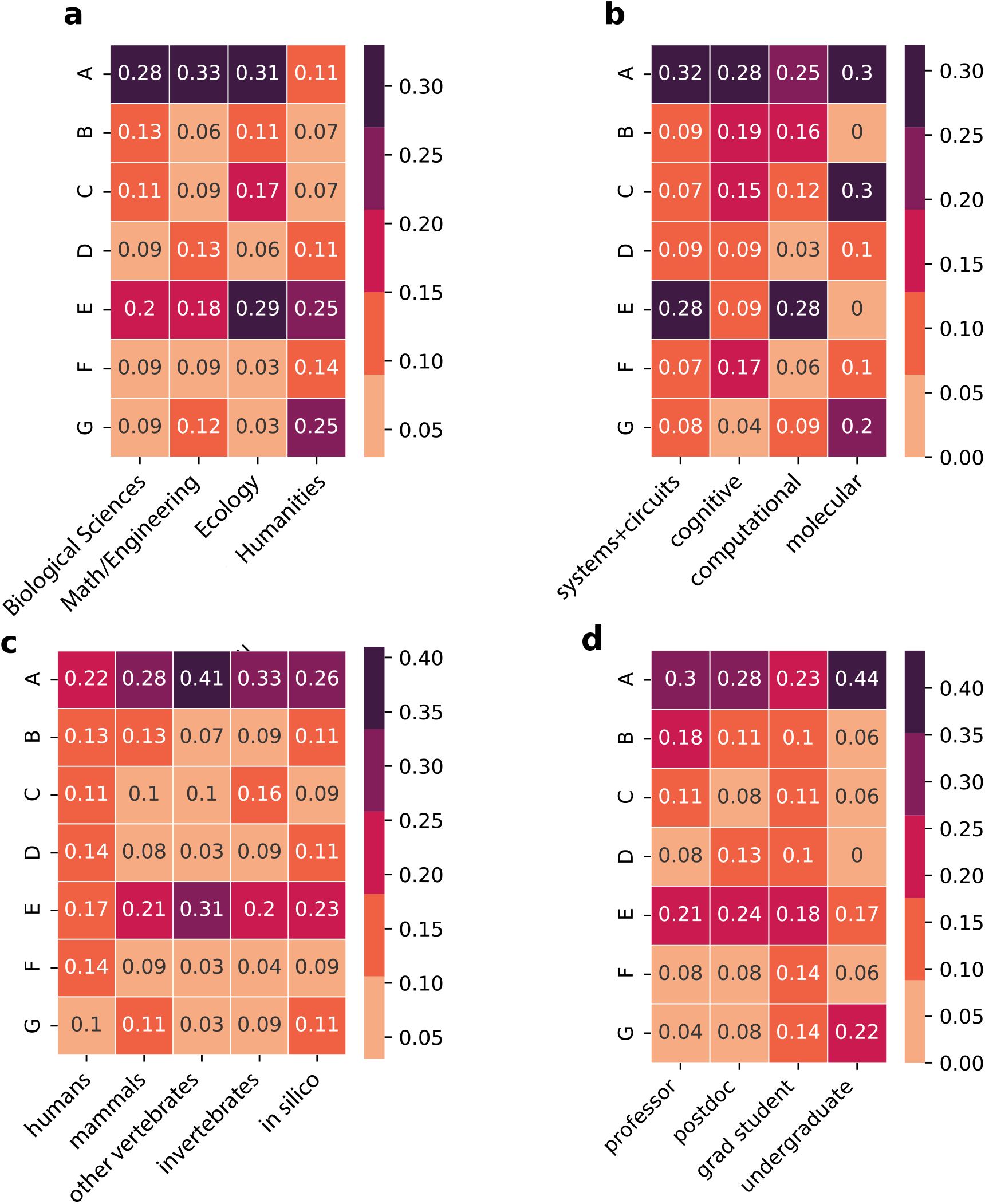
Different groups have distinct distribution of definitions. **a**, Academic fields show different propensities to use each category of ‘behavior’. This is also true of **b** neuroscience specialties, **c** model organism used, **d** and academic seniority. See Methods for definitions of each field.

When broken down by the organism used for the work, those who worked with Humans were the least likely to use ‘broad’ (A) and ‘broad (animals only)’ (E) archetypes but the most likely to use the ‘cognition’ (D) and ‘well-defined/understood behaviors’ (F) definitions. ‘Other vertebrates’ dominated ‘broad’ (A) and ‘broad (animals only)’ (E) and invertebrates were 50% more likely to be a part of ‘broad (but mostly motor’ (C) (Fig 6c).

Undergraduates were predominantly a part of archetypes ‘broad’ (A) and ‘acting with intentions’ (G), whereas professors were the most likely to be part of cluster ‘motor outputs (animals only)’ (B) (Fig 6d).

## Discussion

In this study, we have shown that ‘behavior’ is not universally defined by academics. We have developed and analyzed a survey that has allowed us to identify how the term ‘behavior’ is used, and what it means to people belonging to different academic communities. Using this survey, we have shown that important differences exist in what an academic community would consider to be ‘behavior’, and additionally identified six behavior clusters and seven archetype clusters. These behavioral archetypes correspond roughly to the following labels ‘broad’, ‘animal motor outputs are behavior’, ‘broad but motor’, ‘animal+cognitive’, ‘animal motor+cognitive’, ‘intentional motor behaviors’, and ‘intentional motor outputs by animals’. We speculate that other sub-categories exist - we could, for instance, have cut the tree at other branches or used even more questions and gotten more responses.

Our study is the first report the heterogeneous definitions of ‘behaviors’ across academic communities, as previous studies have focused on a particular sub-field. For example Levitis et al [12] looked at how behavioral biologists define ‘behavior’ and found them to be self-contradictory. However, we have shown that we are able to predict the subjects’ definition of behavior on an individual basis and the definitions are to a great extent self-consistent relative to some underlying set of beliefs. We believe that a follow-up survey focusing on particular terms in behavior, such as ‘decision-making’, ‘foraging’, or ‘computation’ could lead to similar classification of terms that are widely used in literature. It is important to note also that we have taken our utmost care in formulating the questions using precise language but this does not prevent that some of the words might be read in a more complex manner than what we intended to. This is specially true with words that have ambiguous boundaries such as “computer behave”. It might be understood as “it is useful sometimes to consider a computer’s output as a behavior”. We are agnostic to how people interpret these questions and primarily interpret the answers as reflecting some common underlying set of beliefs.

Despite all this, these inconsistent and unclear definitions may obscure otherwise clear differences. For instance, one of the most controversial questions in our survey was Q31: ‘You are wearing a virtual reality headset. You see a virtual tree and reach out to touch it. Is this the same behavior as reaching out to a real tree?’, with roughly equal numbers of Yes, No, and Maybe answers. This has direct relevance to many experiments in neuroscience, where behaviors are ‘simplified’ for experimental tractability: a rodent may be fixed in place and make a ‘decision’ by licking from one of two reward ports. Would this ‘behavior’ be the same in a different context in which the animal was free to explore and move about? it is increasingly apparent that movement itself influences many aspects of the decision process, and animals solve decisions differently when they are head fixed and when they are freely moving [6] - which would not be obviously apparent if we consider them the ‘same behavior’.

Two main schools of thought have dominated the study of behavior: behaviorism, in which behavior can be explained through conditioned motor patterns without regard to cognitive states [17, 18], and cognitivism, in which cognitive processes are thought to be driving behavior and are central to it [9]. More specifically, behaviorism [17, 18] considers behavior as either a reflex to a certain stimulus [5] or one that is shaped by past reinforcement or punishment contingencies. Within the reflex arc conceptual framework [5], behavior is viewed as a set of processes occurring as sequences of distinct steps that are executed in response to perceptual events. Thought is an intermediate step and involves neither sensation nor action.

On the other hand, cognitivism, prevalent in some of the definitions we uncovered in the survey, is a school of thought that emerged in 1950s as a response to the behaviourist school that have neglected study of cognition or saw it as an ephiphenomenon of observable behavior. The cognitivist school believed that cognitive processes seated in the brain, which cannot be directly observed, drive behavior [16] It is crucial to mention that the definitions of behaviors that we unraveled in the survey falls within combinations of the two schools which one would expect as people don’t actually subscribe to these schools explicitly. Nevertheless, these schools give us a taxonomy of the different philosophical lenses through which ‘behavior’ can be viewed and where our subjects would lie - for example if they are more behaviorist than cognitivist.

However, it is unclear if colloquial definitions of behavior actually relate to these definitions. In order to answer these questions, we assigned each of our survey questions to supporting behaviorism, supporting cognitivism, or being unrelated to either (see Methods). Individual responses were assigned a score from −1 to 1 relating to the number of “Yes”es or “No”s that were answered for the relevant questions (Fig 7). We find that very few respondents are strongly Behaviorist or Cognitivist, on either an individual level (Fig 7, histograms) or a group level.

**Figure 7:**
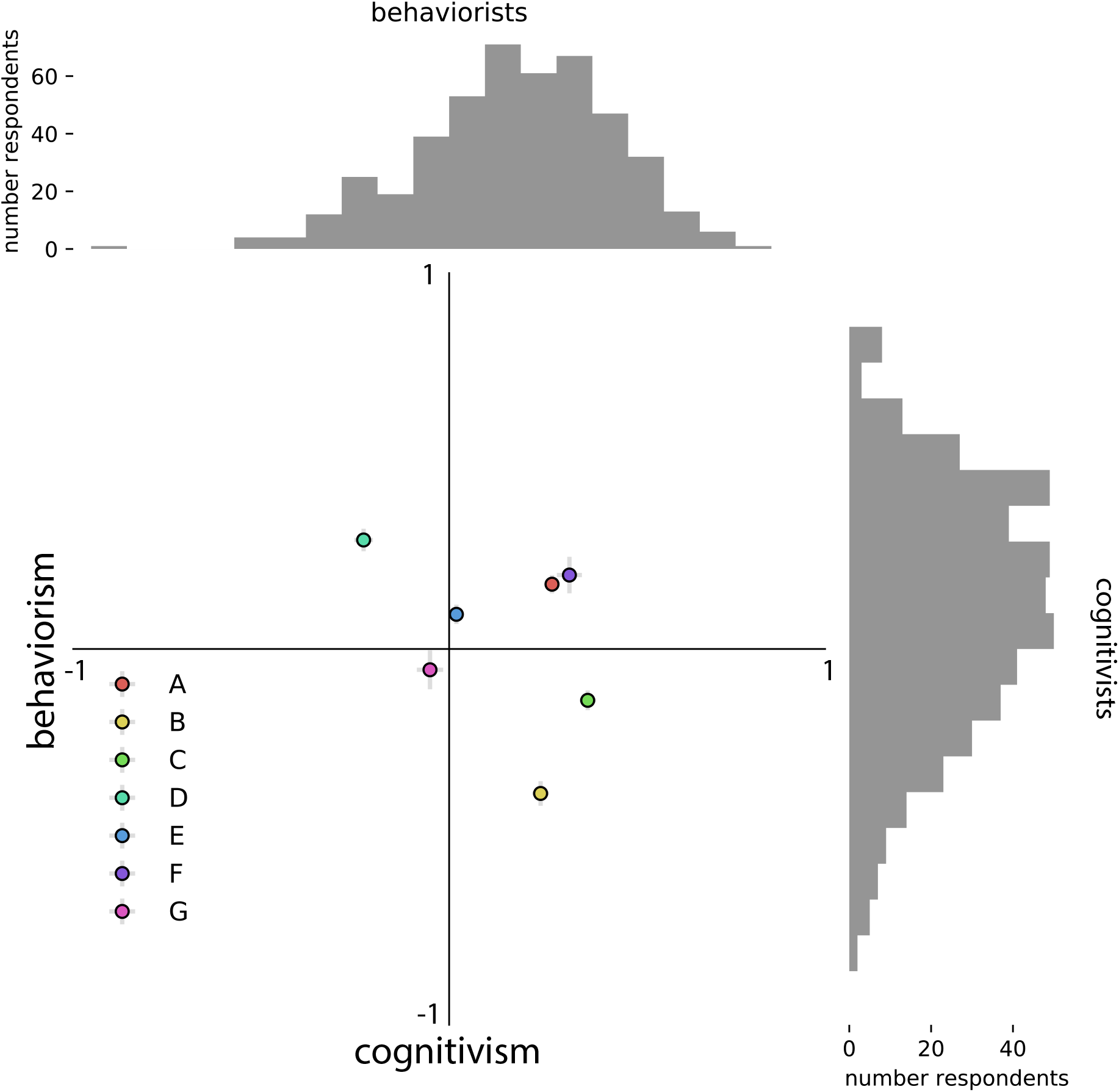
No one cares about behaviorism and cognitivism. Questions were labeled ‘behaviorist’, ‘cognitivist’, or ‘neither’. Responses were scored to either totally agree (+1) or totally disagree (−1) with each category. Respondents did not agree with behaviorism or cognitivism, but used definitions that were a mix of the two. Circles represent mean and error bars are +/− SEM.

Academics often resort to behavioral reductionism by choosing a simplified version of a behavior and then claim that this reflects the entirety of the behavior under study. However, this can be highly misleading for anyone outside of that academic’s field, where they may not understand the implicit definition of the ‘behavior’, with all of its caveats and consequences. This signifies that academic research tends to adopt implicit folk definitions of behavior within a given academic field. For example, a series of nose pokes in an operant chamber may constitute behavior in a systems neuroscience experiment. On the other hand, an anthropologist or a lay person would focus on other aspects, for example the grooming of the animal, the tail movement, etc. We acknowledge the limitations of studying multiple levels of behavioral descriptions simultaneously but we should be cognizant of those limitations, indicating which level we are studying and use more precise language when describing it. This also reflects on the scientific literature that academics publish and if the exact level of description / shortcomings are clarified, inter-disciplinary interactions would be far easier and more fruitful.

We believe this survey makes clear that ‘behavior’ constitutes a wide class of concepts (or, different definitions). One clear suggestion that can be taken from this is the need for integrating different types of ‘behavior’ to make further advances on its study. There are many insightful reviews that have provided a critical analysis of how behavior is studied [8, 10]. To supplement those critiques, here we focus on quantitative analysis of how a sample of academics define behaviors thus providing evidence for the variety of the epistemological frameworks at play in different academic fields. A challenge here is that the hegemony of particular techniques in an academic field can lead to methodological bias which itself delineates the meaning of ‘behavior’. For example in systems neuroscience researchers will systematically use brain recording techniques to look for ‘correlates’ of ‘behavior’, leading to a neurocentric view of behavioral phenomenology. Others will use pure video tracking combined with machine learning algorithms to classify states of behavior regarded in this sense as a behavioralist mechanism. Although we also recognize the difficulty of combining different methods, technological advances are making the possibility of combining multiple techniques more streamlined [14]. It is possible to go further and take an integrative approach which considers how the ecological environment is shaping the behavior along side the brain, the body, and the animal physiological state.

## Acknowledgments

We would like to thank Emily Dennis, Emily Mackevicius, Asif Ghazanfar, Yael Niv, David Barack, Jakob Voigts, Patrick Dylan Rich, Selmaan Chettih, Jess Breda, Sandeep Robert Datta, Benjamin Hayden, and Alex Gomez-Marin for insightful feedback. We would also like to thank Kathleen Quach for the illustrations used in the manuscript.

## Author contributions

A.J.C and A.E. have both designed the survey, analyzed the data and wrote the manuscript

## Code and data availability

Code and data are available at https://github.com/adamjcalhoun/WhatIsBehavior. A version of the survey can be taken at http://adamcalhoun.com/WhatIsBehavior/.

## Methods

### Design

In order to assess whether there is a consistent definition of behavior used across academia, we constructed an online survey using Qualtrics. Questions were developed by the authors who identified potential questions from a wide sample of putative ‘behaviors’, with several drawn from a previous survey performed by other authors [12]. They then sent those questions to colleagues who were experts in animal behavior and ensured that none of the questions were consistently answered ‘Yes’, ‘No’ or ‘Maybe’ in order to identify whether they represented differences of opinion in the field. The survey was then disseminated via Twitter, online mailing lists, as well as to colleagues. Subjects were allowed to answer ‘Yes’, ‘Maybe’, or ‘No’ to all questions. Subjects were asked for metadata consisting of their gender, University or Institute affiliation, country and state of residence, level of seniority, research area, and model system used. Survey questions were randomized and placed in blocks of five questions. Subjects had to answer all the questions before proceeding to the next set of questions. Subjects were free to decline participation in the survey. Subjects were offered the option of leaving their email address to enter a raffle to win a $50 Amazon gift card. The research was approved by the Princeton IRB (IRB# 12895).

### Data analysis

Due to a coding error when designing the survey, subjects did not have to answer the first question in order to continue (’An animal is thirsty. It must choose between two levers, one of which will provide water. Would you refer to the animal pressing the lever as the primary behavior it is producing?’). For consistency, this question was excluded from all analyses. Only data where the questionnaire was finished were used for analysis. Due to the methods used for analysis, we only used responses that finished the entire survey.

### Factor analysis

In order to explore whether answers were similar across fields, data dimensionality was reduced using the python library Prince (available at). Due to the categorical nature of the data, we used Multiple Correspondance Analysis (MCA). Data was transformed into one-hot encodings and then dimensionality was reduced using MCA. Plotting the components (Supp. Fig 2a) showed that the first four components each explain at least 5% of the variance, while subsequent components each explained less than 5% of the variance. Because of this, all analyses were done on the first four components. All data was plotted using the mean and standard deviation across metadata groups.

### Sub-disciplines

Subjects were allowed to enter their ‘sub-field’ in a text box. We categorized these sub-disciplines into four broad disciplines: ‘systems + circuits’, ‘cognitive’, ‘computational’, and ‘molecular’. Subjects were categorized as belonging to ‘systems + circuits’ if they entered ‘systems’, ‘circuit’, or ‘circuits’ in the sub-field box. They were categorized as ‘cognitive’ if they entered ‘cognitive’ or ‘cognition’. They were categorized as ‘computational’ if they entered ‘computational’, ‘theoretical’, or ‘theory’. They were categorized as ‘molecular’ if they entered ‘molecular’ or ‘cellular’. They were categorized as ‘mammals’ if they entered ‘rodents’ or ‘other mammals’, as ‘other vertebrate’ if they entered ‘fish’, ‘birds’, or ‘other non-mammalian vertebrates’, and as ‘invertebrate’ if they entered ‘drosophila’, ‘c. elegans’, or ‘other invertebrates’. ‘Biological sciences’ were ‘neuroscience’, ‘psychology’, ‘biology’, and ‘medicine’, ‘Math/engineering’ included ‘engineering’, ‘statistics’, ‘mathematics’, and ‘machine learning’. ‘Ecology’ included ‘ethology’ and ‘ecology’. Humanities included ‘philosophy’, ‘history’, ‘sociology’, and ‘languages and literature’.

### Hierarchical clustering

Hierarchical clustering was performed using Seaborn and SciPy [22, 24] using the Ward linkage and Euclidean distance (Fig 2).

### Regression analysis of answers

Regression was performed using multinomial logistic regression in scikit-learn [15], with an elastic net penalty and an l1 ratio of 0.5. Answers to questions were fit using 5-fold cross-validation. Questions from the second half of the survey were predicted using one-hot encoding of the answers to questions from the first half, and questions from the first half were predicted from one-hot encoding of the answers to the first half.

### Behaviorism and Cognitivism

After the survey was completed, one author identified questions that a behaviorist or a cognitivist would agree with. For example, Q23 (“A behavior is always potentially measurable”) was chosen to be representative of behaviorism and Q22 (“Does a behavior need to be intentional (does every behavior an animal produces have a purpose)?”) was chosen to be representative of cognitivism.

For each respondent, a belief index was calculated as 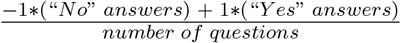.

The following questions were chosen as being behaviorist: Q1, Q4, Q9, Q14, Q15, Q17, Q21, Q23, Q25, Q26, Q27, Q29, Q30, Q33, Q34, Q38, Q40, Q42, Q44, Q46.

The following questions were chosen as being cognitivist: Q0, Q6, Q7, Q8, Q10, Q12, Q13, Q22, Q31, Q28.

This is also available as a CSV file in Supplementary Files and at the GitHub repository.

## Supplemental

**Figure S1:**
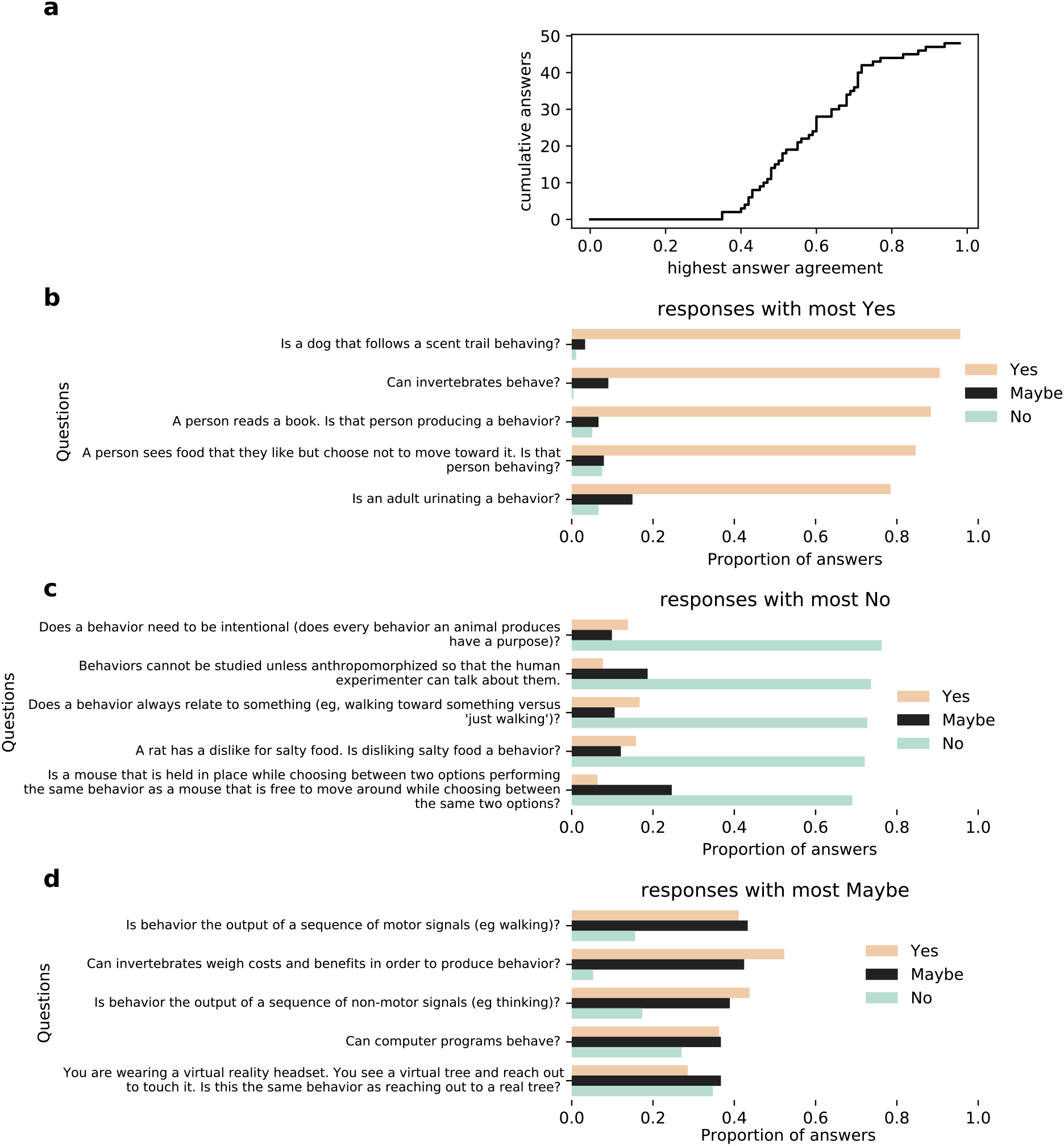
General properties of the responses. **a**, For each question, the most common answer (eg, ‘Yes’) was identified and the probability of that response was plotted. Few questions should ¿80% consensus. **b**, The responses most consistently answered ‘Yes’. **c**, The responses most consistently answered ‘No’. **d**, The responses most consistently answered ‘Maybe’.).

**Figure S2:**
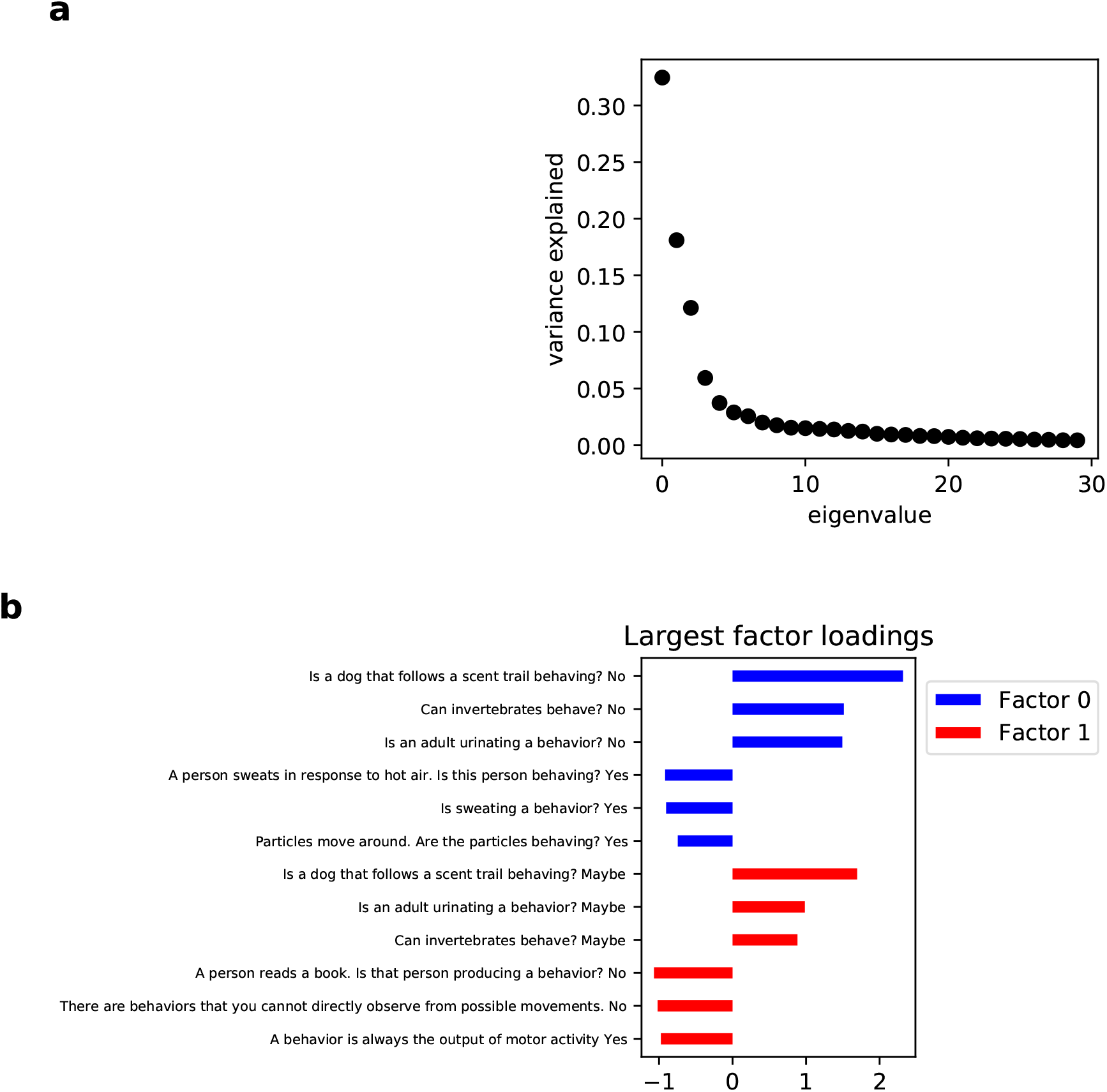
Dimensionality reduction. **a**, The variance explained for each factor of the MCA space. **b**, For the two largest dimensions, the questions that showed the three largest positive and three largest negative loadings for the first (blue) and second (red) factors. Because MCA uses one-hot data, loadings are question-answer pairs.

**Figure S3:**
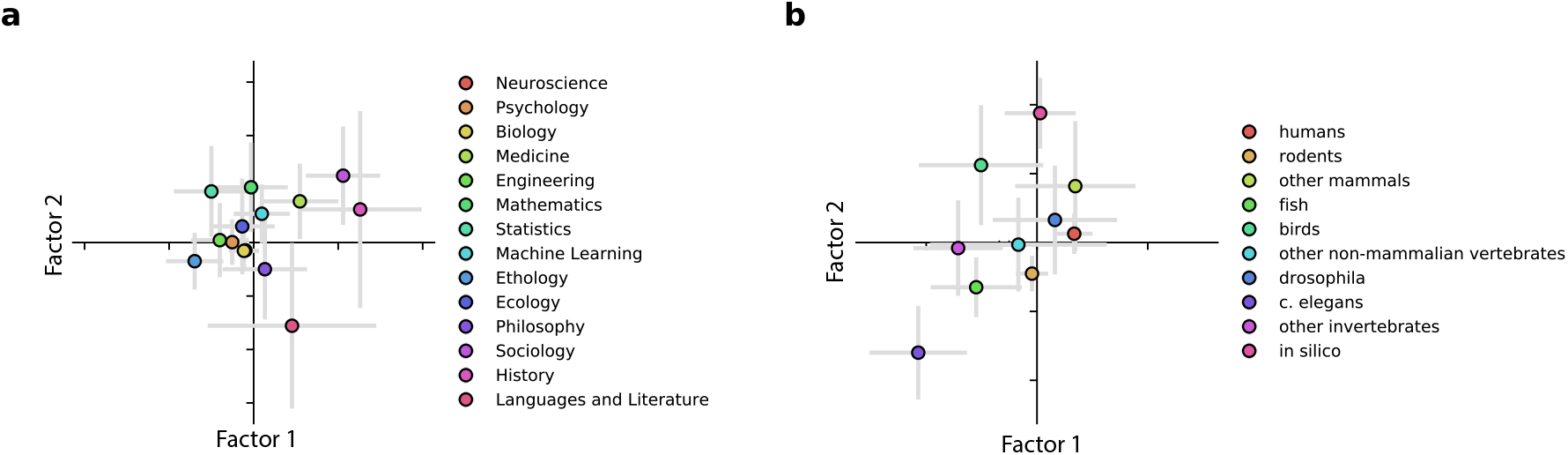
Dimensionality reduction in uncombined categories. The same data as in Figure 1b,d.

**Figure S4:**
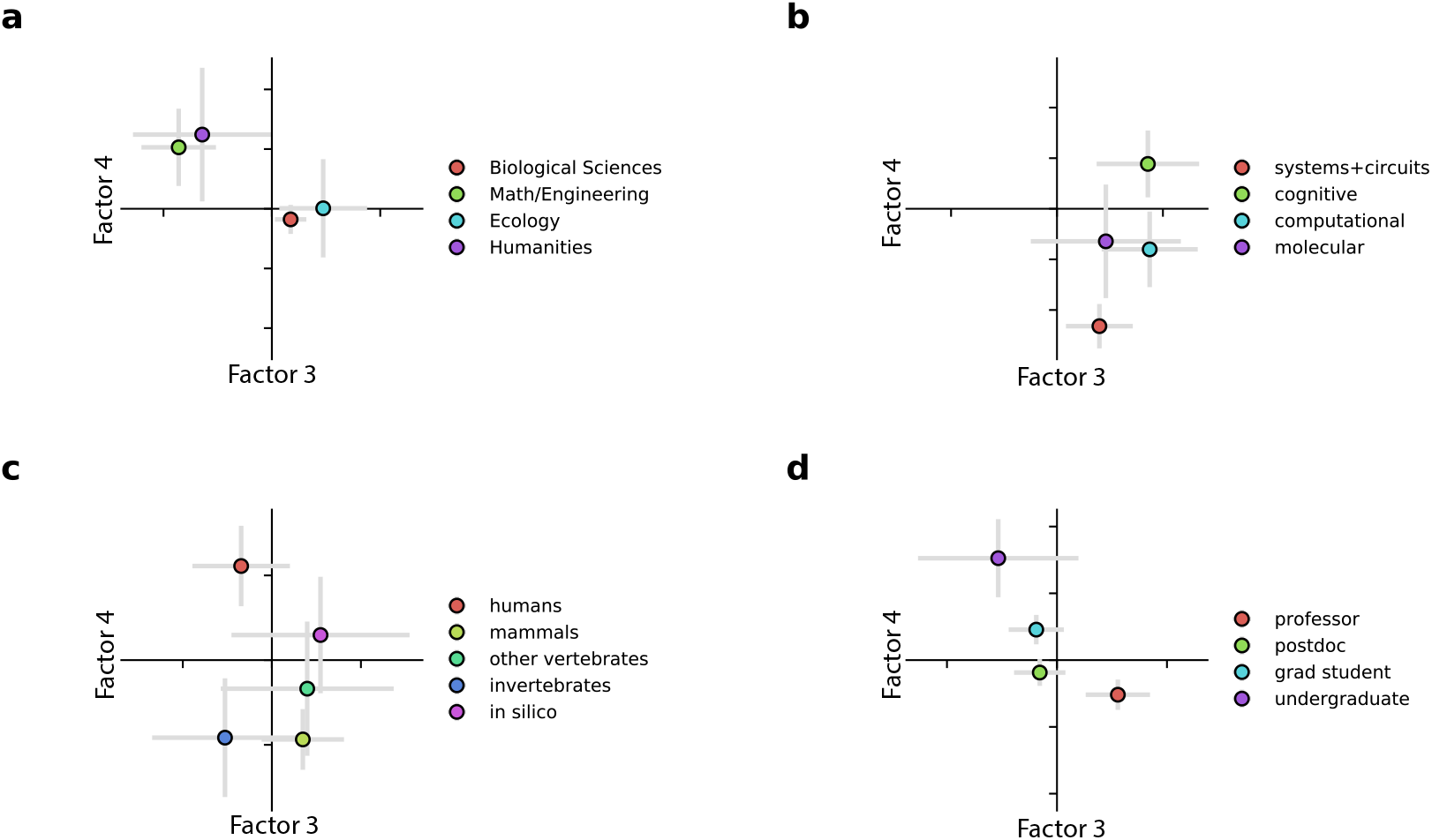
Metadata in higher factors. The same data as in Figure 1b-e, but plotted in the next two largest dimensions.

**Figure S5:**
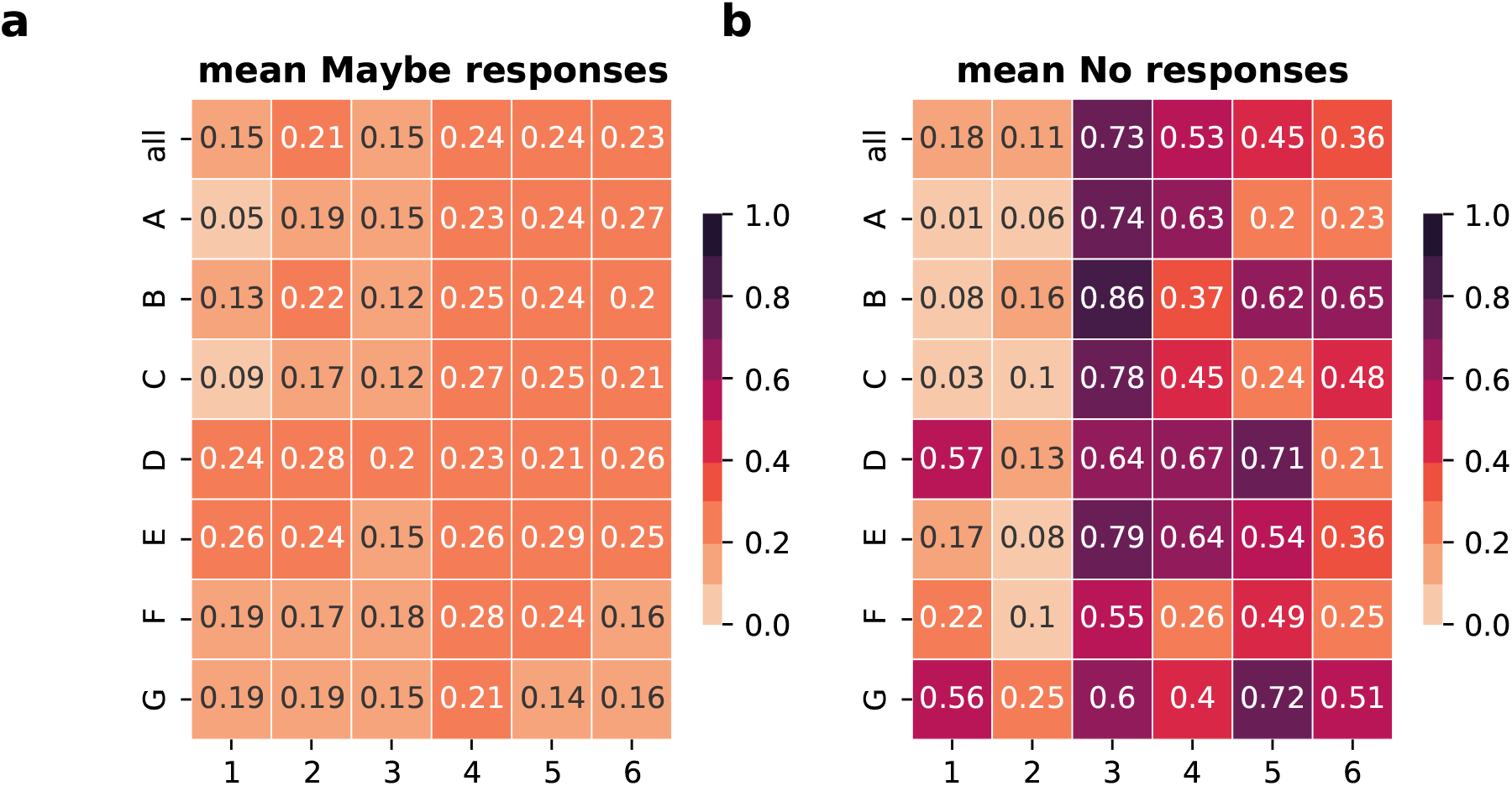
Raw response probabilities. The mean **a**, Maybe responses and **b**, No responses for each behavioral definition and archetype pair. See Figure 3.

**Figure S6:**
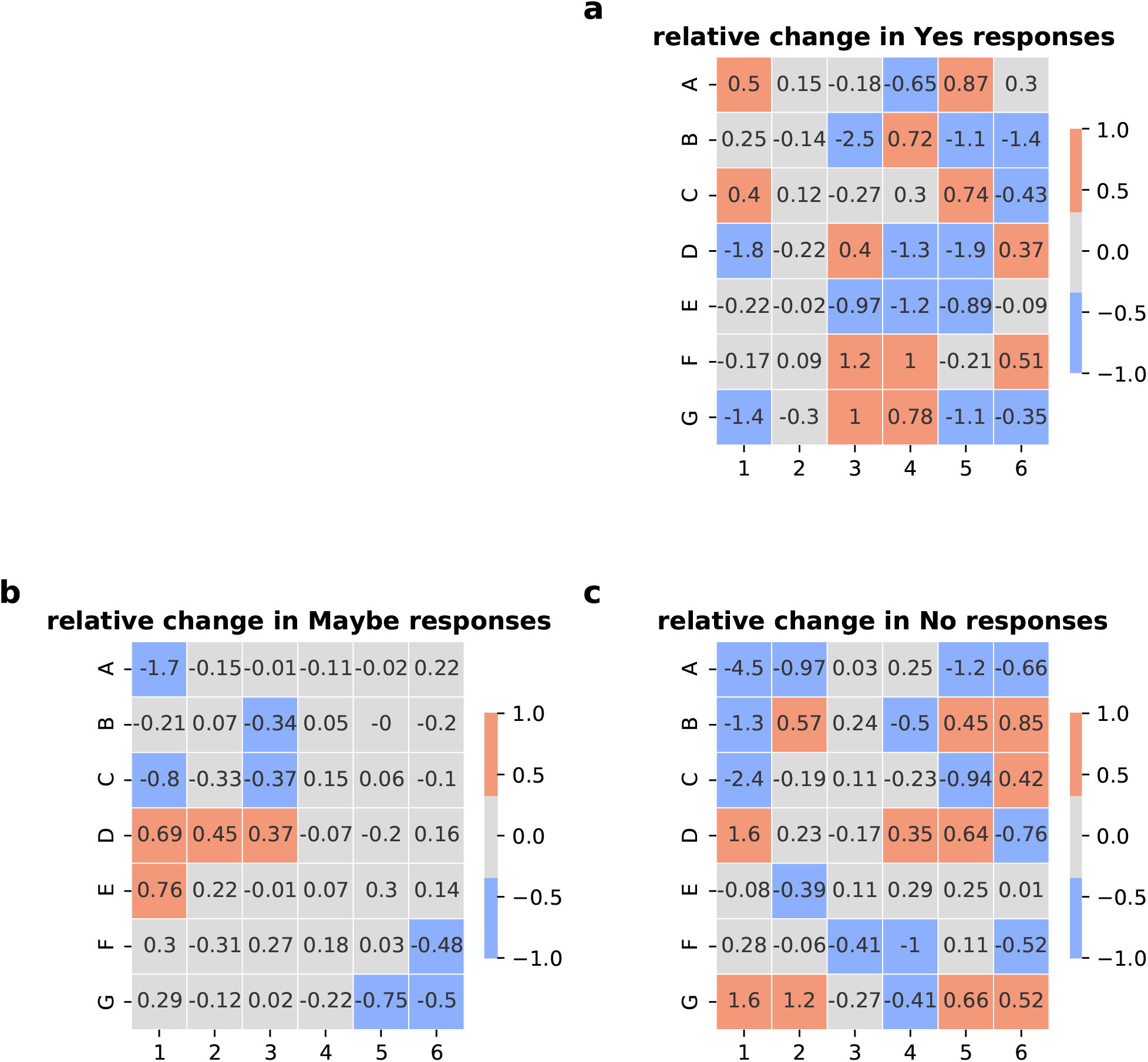
Definition clusters in uncompressed categories. The same data as in Figure 4, replotted by each academic discipline and animal model.

**Figure S7:**
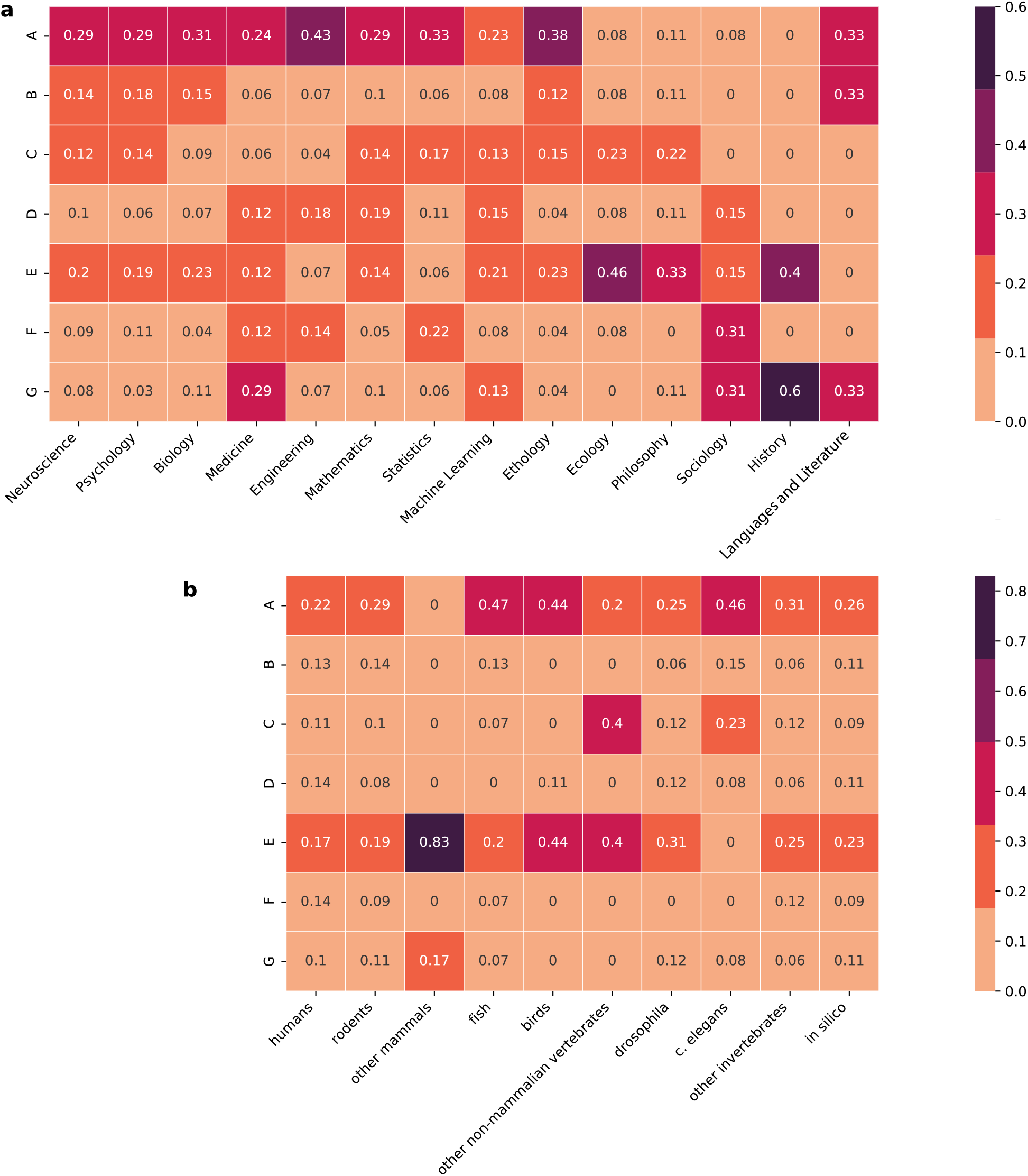
Definition clusters in uncompressed categories. The same data as in Figure 4, replotted by each academic discipline and animal model.

## References

[1] Aristotle. On the soul. 350 BC.

[2] Gordon J Berman, Daniel M Choi, William Bialek, and Joshua W Shaevitz. Mapping the stereotyped behaviour of freely moving fruit flies. Journal of The Royal Society Interface, 11(99):20140672, 2014.

[3] Adam J Calhoun, Jonathan W Pillow, and Mala Murthy. Unsupervised identification of the internal states that shape natural behavior. Nature neuroscience, 22(12):2040–2049, 2019.

[4] David Edward Davis. Integral animal behavior. 1966.

[5] John Dewey. The reflex arc concept in psychology. Psychological review, 3(4):357, 1896.

[6] Benjamin R Eisenreich, Benjamin Y Hayden, and Jan Zimmermann. Macaques are risk-averse in a freely moving foraging task. Scientific reports, 9(1):1–12, 2019.

[7] Molenaar D Fahrenfort JJ Kiverstein JD Seth AK van Gaal S Francken JC, Beereendonk L. An academic survey on theoretical foundations, common assumptions and the current state of the field of consciousness science. PsyArXiv, 2021.

[8] Alex Gomez-Marin and Asif A Ghazanfar. The life of behavior. Neuron, 104(1):25–36, 2019.

[9] John Haugeland. The nature and plausibility of cognitivism. Behavioral and Brain Sciences, 1(2):215–226, 1978.

[10] John W Krakauer, Asif A Ghazanfar, Alex Gomez-Marin, Malcolm A MacIver, and David Poeppel. Neuroscience needs behavior: correcting a reductionist bias. Neuron, 93(3):480–490, 2017.

[11] Brigitte Le Roux and Henry Rouanet. Multiple correspondence analysis, volume 163. Sage, 2010.

[12] Daniel A Levitis, William Z Lidicker Jr, and Glenn Freund. Behavioural biologists do not agree on what constitutes behaviour. Animal behaviour, 78(1):103–110, 2009.

[13] Konrad Lorenz. The foundations of ethology. Springer Science & Business Media, 2013.

[14] Jeffrey E Markowitz, Winthrop F Gillis, Celia C Beron, Shay Q Neufeld, Keiramarie Robertson, Neha D Bhagat, Ralph E Peterson, Emalee Peterson, Minsuk Hyun, Scott W Linderman, et al. The striatum organizes 3d behavior via moment-to-moment action selection. Cell, 174(1):44–58, 2018.

[15] F. Pedregosa, G. Varoquaux, A. Gramfort, V. Michel, B. Thirion, O. Grisel, M. Blondel, P. Prettenhofer, R. Weiss, V. Dubourg, J. Vanderplas, A. Passos, D. Cournapeau, M. Brucher, M. Perrot, and E. Duchesnay. Scikit-learn: Machine learning in Python. Journal of Machine Learning Research, 12:2825–2830, 2011.

[16] Roger Schnaitter. Behaviorism is not cognitive and cognitivism is not behavioral. Behaviorism, pages 1–12, 1987.

[17] BF Skinner. Is it behaviorism? Behavioral and Brain Sciences, 9(4):716–716, 1986.

[18] Burrhus Frederic Skinner. About behaviorism. Vintage, 2011.

[19] Karola Stotz, Paul E Griffiths, and Rob Knight. How biologists conceptualize genes: an empirical study. Studies in History and Philosophy of Science Part C: Studies in History and Philosophy of Biological and Biomedical Sciences, 35(4):647–673, 2004.

[20] Niko Tinbergen. The animal in its world: Explorations of an ethologist, 1932-1972, volume 84. Harvard University Press, 1972.

[21] Uexküll. Umwelt und innenwelt der tiere. Springer Berlin Heidelberg, 1921.

[22] Pauli Virtanen, Ralf Gommers, Travis E. Oliphant, Matt Haberland, Tyler Reddy, David Cournapeau, Evgeni Burovski, Pearu Peterson, Warren Weckesser, Jonathan Bright, Stéfan J. van der Walt, Matthew Brett, Joshua Wilson, K. Jarrod Millman, Nikolay Mayorov, Andrew R. J. Nelson, Eric Jones, Robert Kern, Eric Larson, C J Carey, İlhan Polat, Yu Feng, Eric W. Moore, Jake VanderPlas, Denis Laxalde, Josef Perktold, Robert Cimrman, Ian Henriksen, E. A. Quintero, Charles R. Harris, Anne M. Archibald, Antônio H. Ribeiro, Fabian Pedregosa, Paul van Mulbregt, and SciPy 1.0 Contributors. SciPy 1.0: Fundamental Algorithms for Scientific Computing in Python. Nature Methods, 17:261–272, 2020.

[23] John Von Neumann. The computer and the brain. Yale university press, 2012.

[24] Michael Waskom and the seaborn development team. mwaskom/seaborn, September 2020.

[25] Norbert Wiener. Cybernetics or Control and Communication in the Animal and the Machine. MIT press, 2019.

[26] Alexander B Wiltschko, Matthew J Johnson, Giuliano Iurilli, Ralph E Peterson, Jesse M Katon, Stan L Pashkovski, Victoria E Abraira, Ryan P Adams, and Sandeep Robert Datta. Mapping sub-second structure in mouse behavior. Neuron, 88(6):1121–1135, 2015.

